# Light-driven biological actuators to probe the rheology of 3D microtissues

**DOI:** 10.1101/2022.01.05.475039

**Authors:** Adrien Méry, Artur Ruppel, Jean Revilloud, Martial Balland, Giovanni Cappello, Thomas Boudou

## Abstract

The mechanical properties of biological tissues are key to the regulation of their physical integrity and function. Although the application of external loading or biochemical treatments allows to estimate these properties globally, it remains problematic to assess how such external stimuli compare with internal, cell-generated contractions. Here we engineered 3D microtissues composed of optogenetically-modified fibroblasts encapsulated within collagen. Using light to control the activity of RhoA, a major regulator of cellular contractility, we induced local mechanical perturbation within 3D fibrous microtissues, while tracking in real time microtissue stress and strain. We thus investigated the dynamic regulation of light-induced, local contractions and their spatio-temporal propagation in microtissues. By comparing the evolution of stresses and strains upon stimulation, we demonstrated the potential of our technique for quantifying tissue elasticity and strain propagation, before examining the possibility of using light to create and map local anisotropies in mechanically heterogeneous microtissues. Altogether, our results open an avenue to guide the formation of 3D tissues while non-destructively charting their rheology of 3D tissues in real time, using their own constituting cells as internal actuators.

## Introduction

Tissue engineering holds great potential to develop organotypic in vitro model systems. With their miniaturization in the late 2000’s ^1,2^, such three-dimensional (3D) microtissue models have been used for studying fundamental features of tissue biology such as the repair of wounded fibrous tissue ^3^, or the formation and maturation of myocardial ^4^ and lung tissues ^5^. Their sub-millimetric size has also opened the possibility to use high-throughput, low volume screening of drugs ^6,7^ or functional effects of patient-specific mutations ^8^.

By combining microtissue engineering with magnetic or vacuum actuation, the tissue mechanical properties have recently been assessed and their importance in the tissue formation and physiological function evidenced ^9,10^. For example, Reich et al. demonstrated the respective roles of cell-generated forces and collagen structures in tissue mechanics ^9,11,12^ while Pelling et al. connected remodeling of the cytoskeleton to homeostatic mechanical regulation of tissues ^10,13^. However, using magnetic or vacuum actuations of cantilevers is restricted, by the fixed size of the cantilever, to the characterization of the global mechanical properties of microtissues, and rheological measurements have been shown to be very sensitive to the probe size and the loading history of the mechanical test device used for such characterization ^14,15^. Moreover, such externally applied forces are hardly comparable to cell-generated forces, which constrains their use to study the spatio-temporal regulation of cell forces in tissues. Biochemical treatments, on the other hand, allows the modulation of the signalization pathways responsible for these processes but their inability to target spatially-defined areas of tissues and their low temporal resolution severely limit their potential for further understanding how cell forces are generated, propagated and sensed in physiological and pathological tissues. Indeed, cells are constantly pulling and pushing on their microenvironment, thus assessing, among other things, its mechanical properties ^16^, and probing the mechanical properties of the tissue "from the inside", as the cells naturally do it, would further our understanding of how cells investigate their environment and how cell-generated mechanical signals propagate. Yet, no method currently exists that can locally modulate the contractility of a group of cells within a 3D tissue while simultaneously measuring global tissue contractility as well as both fine-scale cytoskeletal and extracellular architecture.

Thanks to its spatial and temporal resolution, optogenetics has emerged as a powerful tool for spatiotemporally controlling cell signaling ^17^. Using optogenetic control of RhoA activity, several studies recently used light pulses to locally up- or down-regulate cell-generated forces ^18,19^. Valon et al. further demonstrated that these changes in cellular tension were paralleled by tissue deformations in 2D epithelial monolayers ^19^, suggesting an exciting avenue for using cells as mechanical actuators. Such biological actuators could be employed to probe tissue mechanics using light-induced physiological mechanical stimuli.

We combine here 3D microtissues and optogenetics, by engineering 3D microscale constructs of optogenetically-modified fibroblasts embedded within collagen 3D matrices. Using light to control the proximity of RhoA and one of its activator ARHGEF11, we modulate cellular contractility within 3D fibrous microtissues, while microcantilevers report microtissue stress in real time. We demonstrate our ability to control and measure, over space and time, the stress of specific parts of microtissues while simultaneously inferring tissue strain using particle image velocimetry (PIV). We thus investigate the rheology of microtissues, before demonstrating the potential of our technique for quantifying the impact of the rigidity of the cantilever, the extracellular matrix and the differentiation of fibroblasts into myofibroblasts on the tissue elasticity. We then demonstrate the ability of our approach to map local anisotropies in mechanically heterogeneous microtissues, as well as to influence tissue architecture using repetitive stimulations during tissue formation. Together, these results highlight a unique approach to examine the effects of various parameters such as mechanical preload, tissue maturation, matrix architecture and stiffness on both the ability to propagate physiological mechanical signals and the dynamic mechanical properties of engineered fibrous microtissues.

## Results

### Optogenetic stimulation of microtissues

We used optogenetics to generate contractions in parts of engineered microtissues (Fig. 1.A-B). Our strategy was to trigger the activation of the small GTPase RhoA, a major regulator of cellular contraction ^20^. To this end, we used NIH3T3 cells stably expressing a Cry2-CIBN optogenetic probe (opto-RhoA fibroblasts) to dynamically control with blue light the localization of ArhGEF11, an upstream regulator of RhoA ^21^. As previously described, ArhGEF11 was recruited to the cell membrane upon blue light stimulation, thus activating RhoA and subsequently cell contractility (Fig. 1.C)^19^. We embedded these opto-RhoA fibroblasts in a neutralized collagen I solution within microfabricated PDMS wells containing two T-shaped microcantilevers. Over time of cultivation, the fibroblasts spread inside the collagen and spontaneously compacted the matrix to form opto-RhoA microtissues that spanned across the top of the pair of cantilevers (Fig. 1.A), which deflection was measured to quantify tissue tension ^2^. During tissue formation, a baseline static tension developed due to compaction of the gel by the collective action of the fibroblasts. Once the tissue formed, we used a digital micro-mirror device (DMD) to illuminate parts of these opto-RhoA microtissues. To demonstrate our ability to apply local stimulations, we consecutively illuminated the left and right halves of microtissues while measuring the microtissue tension as well as the local displacements using PIV (Fig. 1.D-E, Supp. movie 1). Immediately after a pulse of light (characterized by an irradiance of 1.7 mW/mm^2^ and a duration of 500 ms, i.e. an energy of 0.9 mJ/mm^2^), the tissue in the stimulated area contracted, inducing a quick increase in tissue tension (half-time to maximum increase 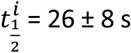), before slowly relaxing back to baseline (half-time of decrease from maximum 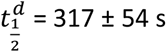) (Fig. 1.D-F). Fig. 1.F shows the tension increase corresponding to the local contractions of the left and right halves of the microtissues. For both stimulations patterns, the microtissue tension increased to a similar level and decreased back to baseline level, thus indicating an elastic response without plastic deformation. By engineering microtissues composed of wild type (WT) NIH-3T3 fibroblasts, we confirmed these light induced contractions were due to the optogenetic construction and inexistent in WT microtissues (Fig. 1.D-F). Of note, opto-RhoA microtissues were broader and generated more tension than WT microtissues, but both generated similar baseline stress σ_xx_ (i.e. tissue tension obtained from the cantilever deflection divided by the cross-sectional area in the center of the tissue) (Fig. 1.G). Furthermore, the local displacements were mostly directed along the longitudinal axis, suggesting a strongly anisotropic contraction. To further investigate the geometry of deformation, we next stimulated a discoidal area of microtissues suspended between two and four cantilevers.

**Figure 1.**
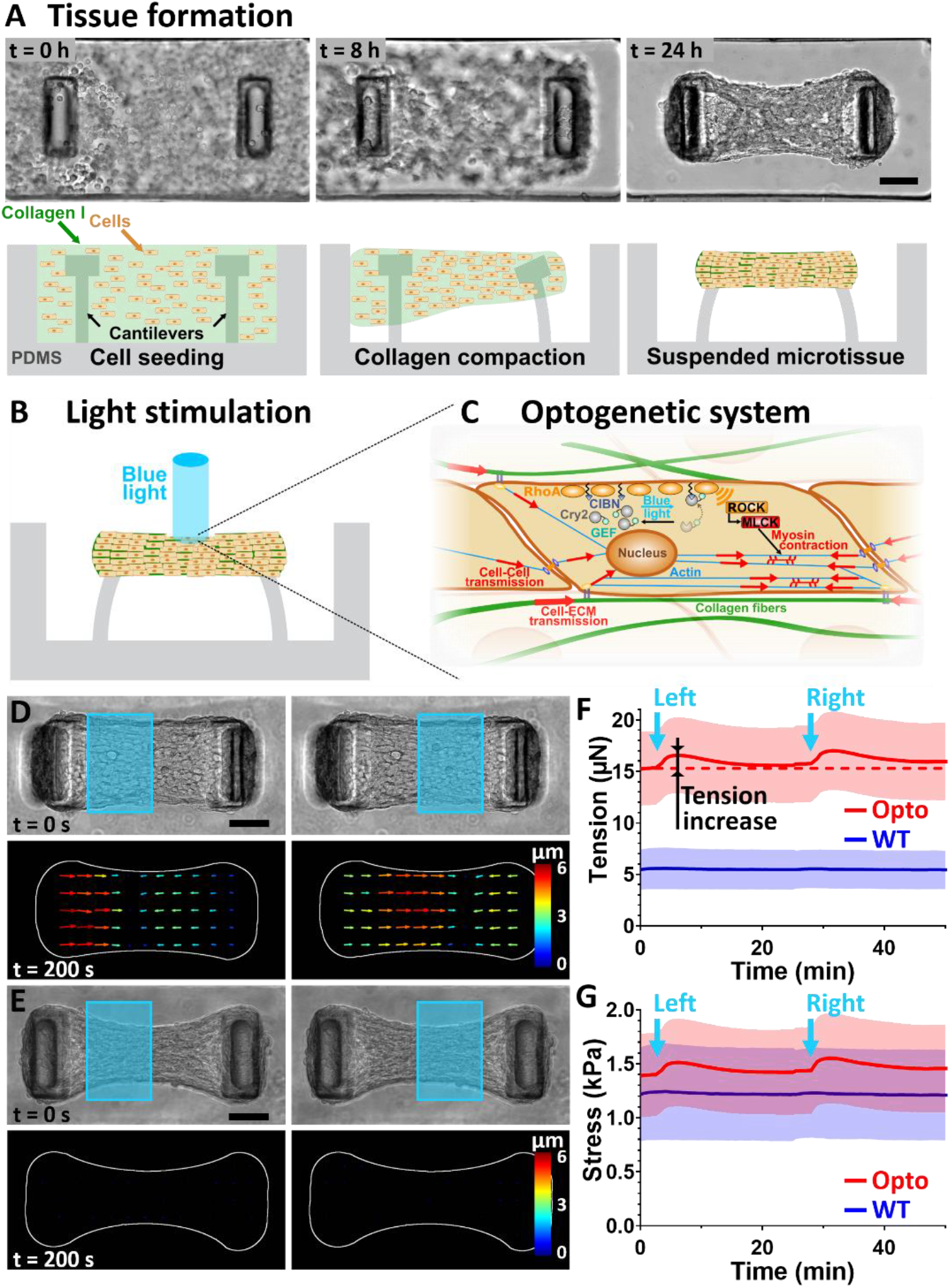
Light-controlled contraction of microtissues. (A) Representative top view images and side view schemes depicting the formation of a microtissue. Over the first 24 h of cultivation, the fibroblasts spread and spontaneously compact the collagen matrix. The two T-shaped cantilevers anchor and constrain the contraction of the collagen matrix to form a microtissue that spans across the top of the pair of cantilevers. (B) Schematic illustrating the light-stimulation of a microtissue composed of Opto-RhoA fibroblasts. (C) Schematic of the ARHGEF11(DHPH)-CRY2/CIBN-CAAX optogenetic system to control cellular contractility. Upon blue light stimulation, CRY2 changes conformation and binds to CIBN, bringing the RhoA-activator ARHGEF11 in close proximity of RhoA, which in turn induces a contractility increase. (D) Upon blue light illumination of its left- or right-half, an opto-RhoA microtissue contracts locally, as shown by the PIV-tracking of the displacements, whereas a WT microtissue is unaffected (E). For readability reasons, only half of the vectors are represented. Scale bar is 100μm. Temporal evolution of the tension (F) and stress *σ_xx_* (G) generated by Opto-RhoA or WT microtissues upon the simulation of their left- and right-half. Data are the average of 20 microtissues ± SD.

### The optogenetically-induced contraction follows the anisotropy of the microtissue architecture

A major advantage of optogenetics over drug or genetic approaches is the possibility to stimulate spatially defined areas of a tissue of interest. We thus investigated the mechanical response of opto-microtissues of which only a small area is stimulated with blue-light. We stimulated a 50 μm diameter disc in the center of the microtissue (Fig. 2.A, Supp. movie 2) and mapped the resulting displacements over the whole microtissue by using particle image velocimetry (PIV). We observed that the stimulated central region barely moved upon illumination while the left and right parts of the microtissues were displaced inwards, toward each other, indicating compaction of the stimulated area (Fig. 2.B). Despite a discoidal light stimulation, the resulting displacements were strongly polarized, with a mean angle of 16 ± 7 ° and more than 80% of the displacements presenting a lower than 30° angle with the longitudinal x-axis of the tissue (Fig. 2.C). This polarized displacement field corresponds to a strongly anisotropic field of deformation, as the x-component of the strain (*ε_xx_*) was more than 15-times larger than its y-component (*ε_yy_*) (Fig. 2.D-F, Supp. movie 2). We defined the anisotropy coefficient (AC) as the difference between *ε_xx_* and *ε_yy_*, normalized by their sum. In the case of these microtissues suspended between two cantilevers, we measured an AC of 1.2 ± 0.2, indicating a strong anisotropy along the x-axis. This quasi-1D response suggests a strong anisotropy of either the cytoskeleton of the constituting cells or the surrounding matrix. We simultaneously stained actin and collagen in microtissues and quantified their alignment. The confocal images showed a compacted collagen core, sparsely populated with fibroblasts, surrounded by a highly cellularized peripheral shell (Supp. movie 3), consistent with previous observations ^9,22^. We also found that the longitudinal contraction induced by light correlated with a strongly anisotropic architecture of both actin and collagen, with more than 80 % of the actin fibers and more than 70 % of the collagen fibers aligned within less than 30 ° of the x-axis (Fig. 2.G-L, Supp. movie 3). As cortical actin is prominent, cell alignment correlates with actin alignment, which is coherent with the fact that fibroblasts line up and remodel their extracellular matrix to align with the principal maximum strains developed during tissue formation ^23–25^. Furthermore, we show here using optogenetics that any additional contraction, even local, is constrained by the anisotropic conformation of the tissue.

**Figure 2.**
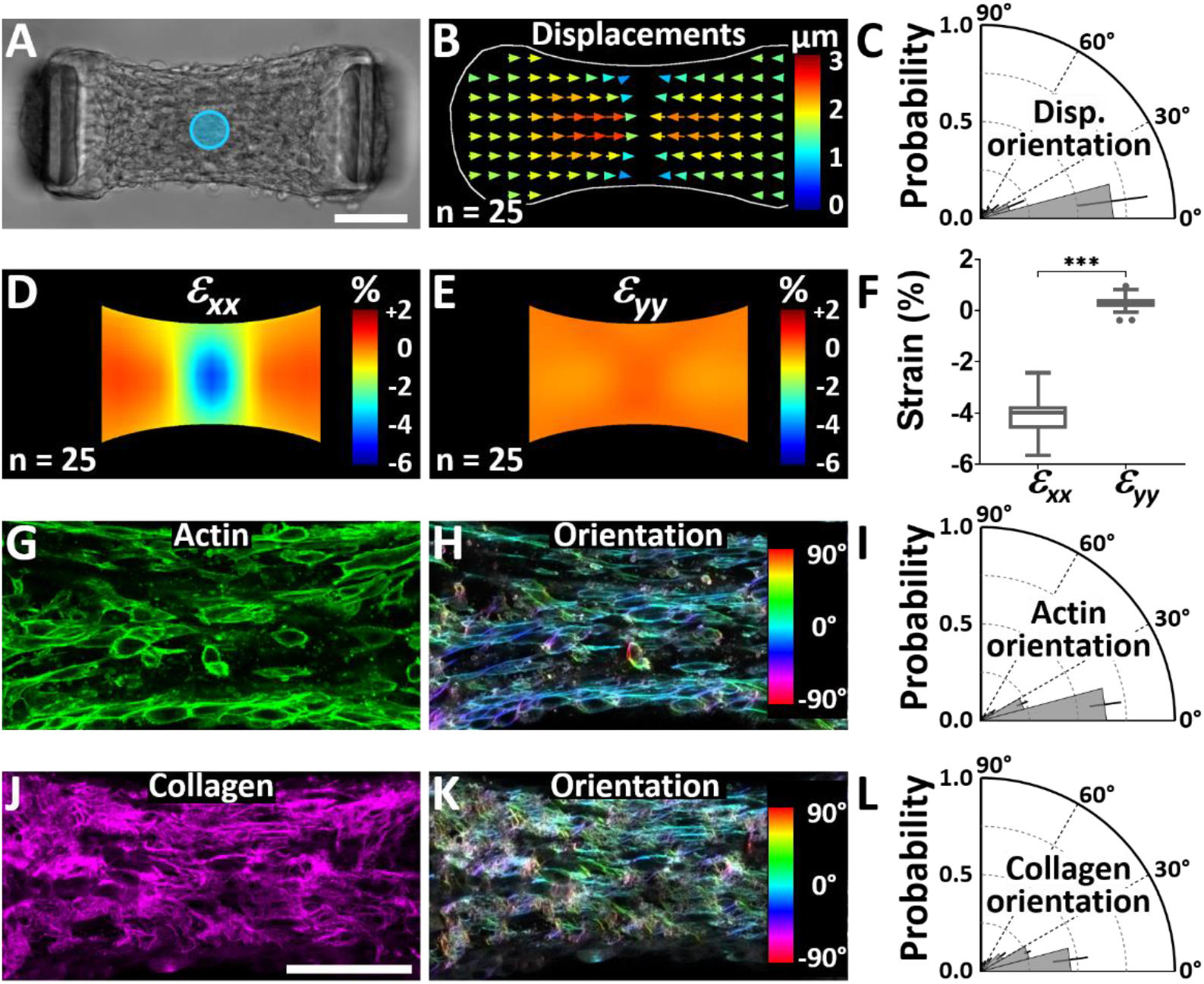
The anisotropy of the optogenetically-induced contraction correlates with the microtissue architecture. (A) Representative microtissue where the blue disc represents the 50 μm diameter isotropic stimulation area. (B) Resulting average displacement field (for readability reasons, only half of the vectors are represented) and (C) polar plot of its orientation. Data are presented as mean ± SD with n = 25 microtissues. Resulting average strain fields (*ε_xx_* in D, *ε_yy_* in E) and comparison of the average strain amplitudes in the area of stimulation (F). Data are presented as Tukey box plots with n = 25 microtissues. ***p < 0.001 between *ε_xx_* and *ε_yy_* (G) Confocal slice of the fluorescent staining of actin in a representative microtissue and (H) corresponding color-coded map of the actin orientation. (I) Polar plot showing the anisotropic orientation of actin fibers along the x-axis. (J) Confocal slice of the fluorescent staining of collagen in a representative microtissue and (K) corresponding color-coded map of the collagen orientation. (L) Polar plot showing the anisotropic orientation of collagen fibers along the x-axis. Data of actin and collagen orientations are presented as mean ± SD with n = 27 microtissues. Scale bars are 100 μm.

### Dynamic of optogenetically-induced strains

Having shown the quasi-1D deformation of anisotropic microtissues spanning between two cantilevers, we next sought to further investigate the spatio-temporal strain patterns induced by such local, optogenetically-induced contractions. We stimulated the left half of microtissues and used PIV to map the resulting displacements and infer the corresponding strain *ε_xx_* (Fig. 3, Supp. movie 4). We observed that the stimulated half left of the microtissue was compressed, up to 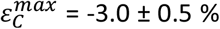, while the non-stimulated half right of the tissue was stretched, up to 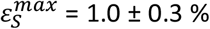. This result indicates that when stimulated cells contract, compact and compress their surroundings. This compression is equilibrated by both the stretch of the non-stimulated part of the tissue and the deflection of the micropilllars. Figure 3. C-D shows the evolution of tissue stress *σ_xx_*, compression and stretch over time, indicating two time delays *τ_C_* = 70 ± 39 *s* between the maximum stress and the maximum compression, and *τ_S_* = 200 ± 64 *s* between the maximum stress and the maximum stretch. By stimulating only one part of the microtissue, we were thus able to simultaneously measure both the stress *σ_xx_* and the strain *ε_xx_*, inferred from the PIV tracking of local displacements (Fig. 3. E). These results suggest the possibility to use optogenetic stimulation for assessing the mechanical properties of the microtissue.

**Figure 3.**
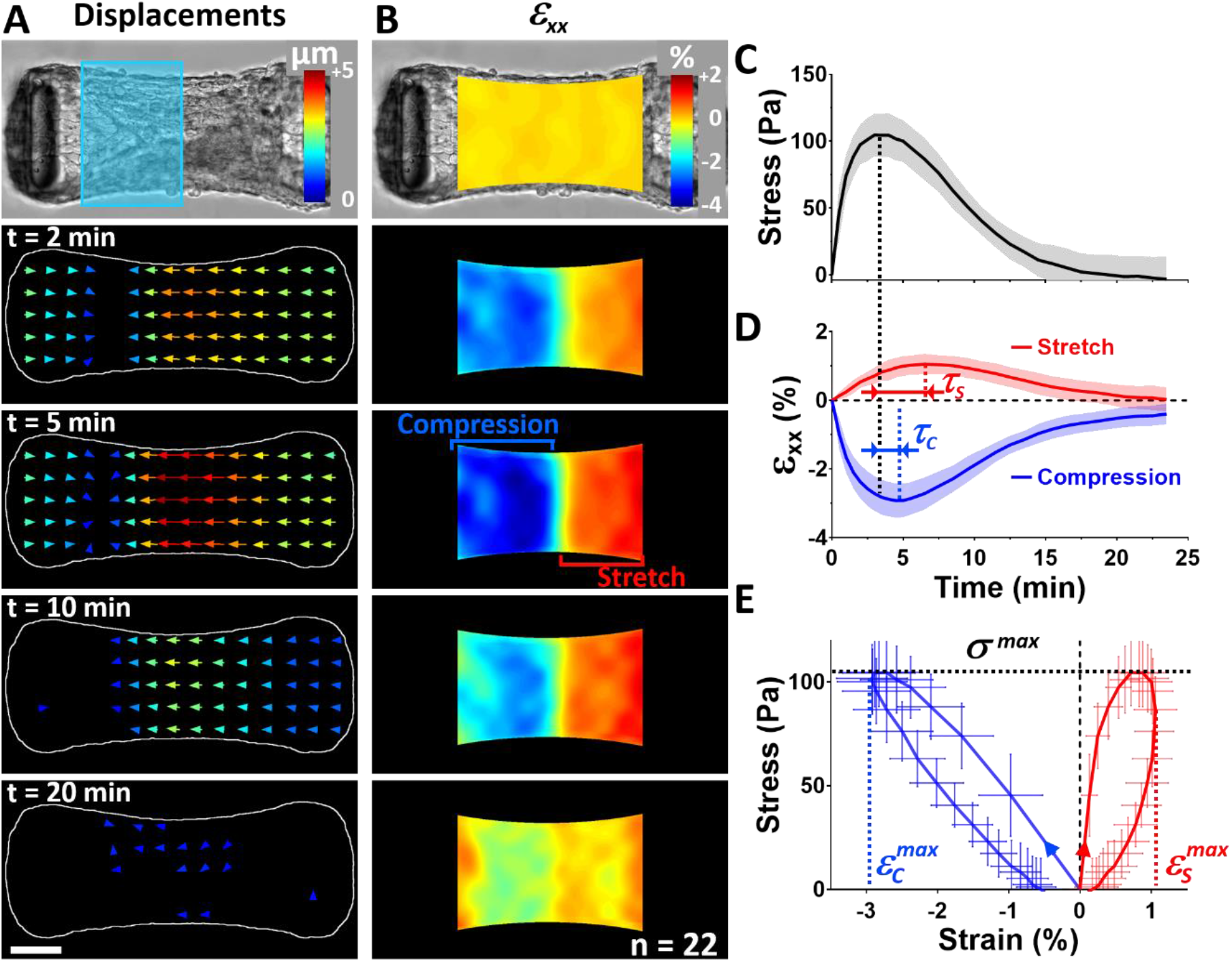
Light-induced local contractions evidence the viscoelastic properties of microtissues. Temporal evolution of the average displacement (A) and *ε_xx_* strain field (B) after stimulation of the left half of the tissue by blue light (symbolized by a blue rectangle in A). (C) Evolution of the stress *σ_xx_* over time and (D) resulting strain *ε_xx_* decomposed in its positive (stretch, in red) and negative (compression, in blue) components. (E) Corresponding stress-stretch (in red) and stress-compression (in blue) curves. The dotted lines indicate the maximum stress *σ^max^* (in black), the maximum stretch 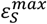 (in red) and the maximum compression 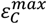 (in blue). Data are the average of 22 microtissues ± SD.

### Inferring mechanical properties of stretched areas

In order to probe the mechanical properties of the microtissue, we used the stimulated half of the tissue as a mechanical actuator stretching the non-stimulated half. We first compared the amplitudes of stress and stretch while varying the width of the stimulated area from 20 to 100 μm (Fig. 4, Supp. movie 5). We thus demonstrated that the force increase was proportional to the area of the stimulated zones (Fig. 4.A and Supp. Fig. 1.A-B). As the cell density is roughly homogenous along the whole length of the microtissue (Supp. Fig. 1.C-H), this result indicates that the light-induced contraction is directly proportional to the number of stimulated cells. As the contraction is quasi-1D, the cross-section of the microtissue is unchanged when the width of stimulation is varied, and the resulting, optogenetically-induced stress increase is also proportional to the width of stimulation, ranging from 98.5 ± 11.2 Pa to 223.9 ± 32.7 Pa when the stimulation width increases from 20 to 100 μm, respectively (Fig. 4.C).

As a result, the average stretch of the non-stimulated part increased from 0.6 ± 0.2 % to 1.2 ± 0.3 % for the same range of stimulation width (Fig. 4.B-C, Supp. movie 5). Upon actuation by the stimulated part, the non-stimulated part of the tissue thus exhibits viscoelastic characteristics when undergoing deformation. The maximum stretch 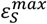 is linearly depending on the maximum applied stress *σ_max_*, although peaking with a delay *τ_S_*, and both stress and *ε_xx_* strain returns to 0 after stimulation. Consequently, we defined the apparent elastic modulus of the tissue *E* as the maximum stress *σ^max^* divided by the maximum stretch 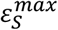 and observed that *E* was logically independent of the stimulation width, with an average value of 19.2 ± 5.0 kPa (Fig. 4.D). We did not measure any significant difference in the time delay *τ_S_* between maximum stress and stretch when varying the size of the stretched part (Fig. 4.D), thus demonstrating that this delay is not a poroelastic effect occurring through the redistribution of the fluid within the microtissue but rather to the relaxation of the actomyosin cytoskeleton.

**Figure 4.**
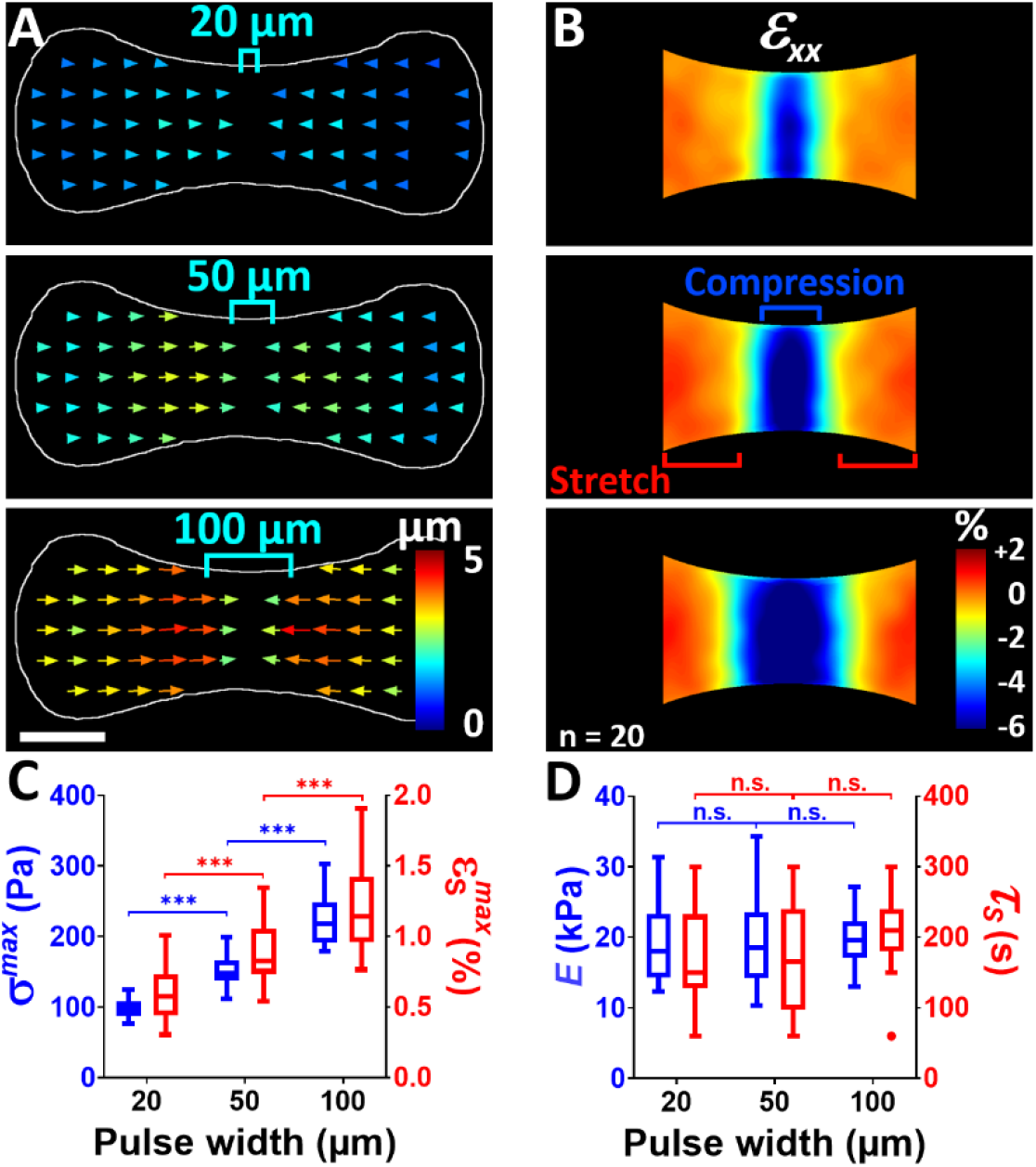
Light-driven probing of the apparent elastic modulus of microtissues. Average displacement (A) and *ε_xx_* strain (B) fields resulting from 20 μm, 50 μm and 100 μm wide light stimulations (symbolized by blue brackets). For readability reasons, only half of the displacement vectors are represented. Scale bar is 100 μm. (C) Corresponding maximum stress *σ^max^* and stretch 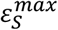. (D) Resulting elastic modulus *E*, reported as 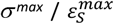, and time delay *τ_S_* between reported as *σ^max^* and 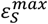 in function of the pulse width. Data are presented as Tukey box plots with n = 20 microtissues. ***p < 0.001 between conditions. n.s. stands for non-significant (i.e. p > 0.05).

Of note, although the range of possible frequencies is limited compared to external actuators, the stress-strain curves we obtained could be analyzed with rheological models commonly used for collagenous microtissues (Supp. Fig. 2), such as the Kelvin-Voigt model (a spring and a dashpot in parallel), the standard linear solid (SLS) model (a spring and a dashpot in series, in parallel to another spring), or the stretched exponential (SE) model (a sum of Maxwell bodies, i.e. a spring and a dashpot in series, with a specific distribution of time constants) ^13,26^. Fitting such models to the data presented in Figure 4 for a stimulation width of 50 μm lead to consistent relaxation times τ between 101 and 135 s for the three different models. Although the Kelvin Voigt model captured the overall shape of the mechanical response and gave an elastic modulus of 18.7 kPa close to our experimental measurements, it failed to capture both the short- and long-time behavior. The SLS and the SE models lead to similar elastic constants (E_1_ = 56.3 kPa and E_2_ = 14.7 kPa for the SLS model, E_1_ = 65.0 kPa and E_2_ = 10.0 kPa for the SE model), but the SE model allowed for better fitting the long-time recovery response, thanks to the dimensionless constant β = 0.66 that captures a specific distribution of timescales (i.e. when β = 1, the SE model behaves as a SLS model, whereas when β decreases, the distribution of timescales broadens). Such distribution is similar to previously obtained results ^13^ and describes the broad distribution of inter-related timescales inherent to the viscoelastic heterogeneities of the different microtissue components.

We then tested the robustness of our approach by assessing the rheological properties of microtissues under different conditions (Supp. Fig. 3). We first investigated the influence of the boundary conditions on the rheological properties of the microtissue. To this end, we stimulated locally microtissues suspended between cantilevers presenting a spring constant between 0.20 to 1.12 N/m, which have been previously shown to influence tissue contractility and stiffness ^2,4,9^. We measured the resulting stretch of the non-stimulated zones and inferred elastic moduli and temporal delay between maximum stress and stretch. In good agreement with previous measurements using various spring constants ^9^, we found that the elastic modulus increased from 6.2 ± 2.5 to 21.4 ± 8.3 kPa with increasing cantilever spring constant (Supp. Fig. 3).

We also observed, for the first time to our knowledge, that the delay *τS* between maximum stress and stretch more than doubled, from 133 ± 54 s to 282 ± 75 s, when the cantilever spring constant was increased from 0.20 to 1.12 N/m. As cellular contractility is the main regulator of tissue tension, while collagen architecture was shown to impact predominantly the tissue stiffness ^9,11,12,26^, our results suggest that this delay *τ_S_* is due to the rupturing and reforming of bonds within the cytoskeleton (i.e. between actin filaments, between actin and myosin or between cadherin and catenin), in coherence with the previously shown stretch-induced perturbation of the cytoskeleton ^13,27–29^. To test this hypothesis, we interrogated the dependence of the measured elastic modulus and time delay on the amplitude of the light-induced contraction. As the photo-activation of CRY2 depends on both the amplitude and the frequency of the light pulses ^21,30^, we varied the amplitude of contraction by varying either the irradiance of a single light pulse, or the number of 1-minute-spaced light pulses (Supp. Fig. 4). We measured no significant differences in elastic modulus, while the delay *τ_S_* between maximum stress and *ε_xx_* strain increased with the amplitude of contraction. We further verified that the light-induced contraction was not inducing plastic, irreversible deformation of the collagen and/or actomyosin network. To this end, we illuminated the left half of microtissues with six successive light stimulations, spaced far enough apart in time to allow complete relaxation. We did not measure any difference in elastic modulus or time delay between the different stimuli (Supp. Fig. 5). Altogether, these results demonstrate that using the cells themselves as mechanical actuators allows to probe, as the cells naturally do, reversibly and non-destructively, the elasticity of fibrous microtissues. Moreover, this approach evidences the stress-dependent dynamic of strain propagation in such tissues.

**Figure 5.**
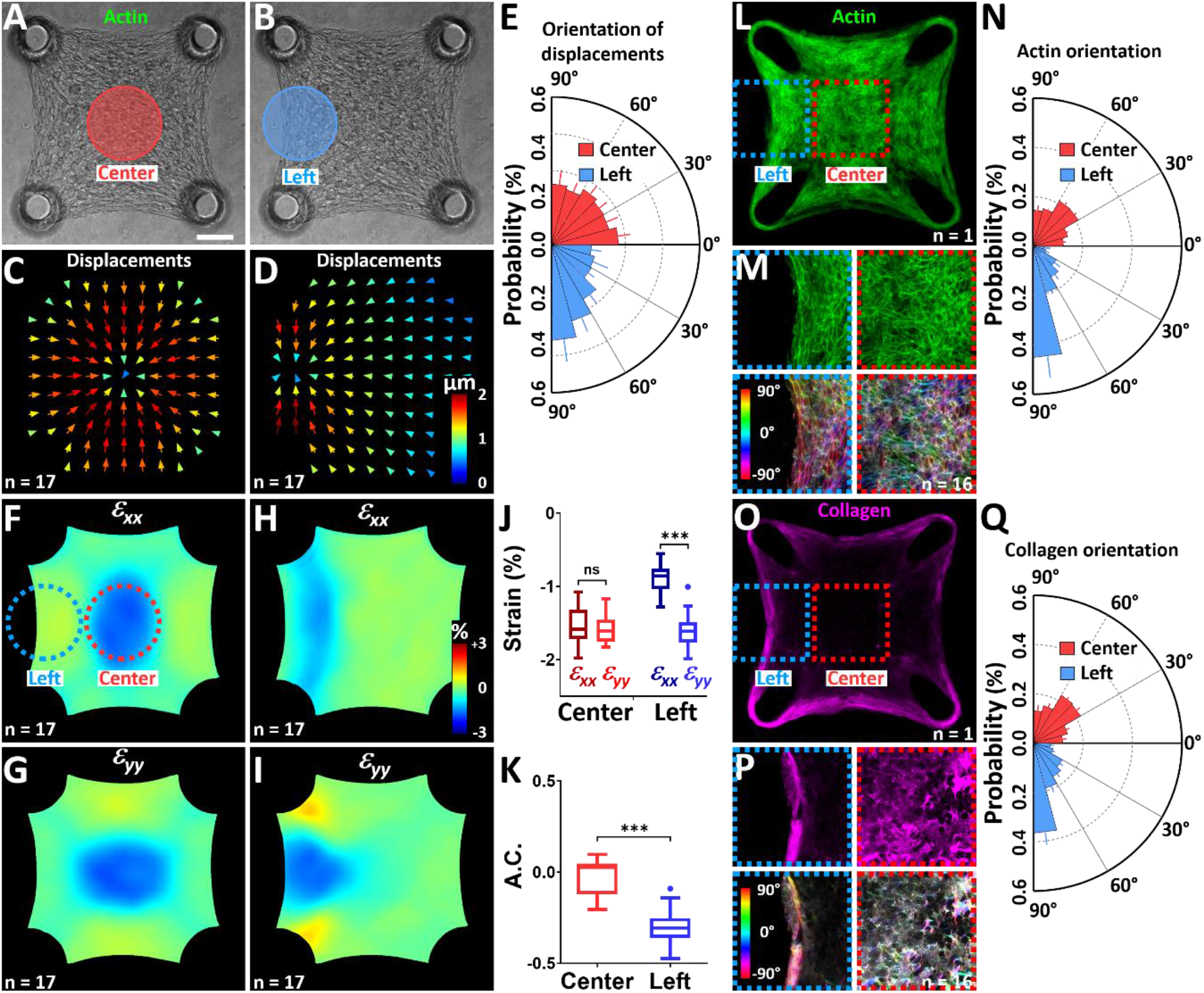
Optogenetically-mapped mechanical heterogeneities correlate with actin and collagen architecture. (A) Representative square microtissue where the red (A) and blue (B) discs represent the centered and eccentric 200 μm diameter isotropic stimulation areas, respectively. Corresponding average displacement fields for the centered (C) and eccentric (D) stimulations. For readability reasons, only half of the vectors are represented. (E) Polar plot of the orientations of the corresponding displacement vectors. Data are presented as mean ± SD with n = 17 microtissues. Resulting average strain fields for centered (*ε_xx_* in F, *ε_yy_* in G) and eccentric (*ε_xx_* in G, *ε_yy_* in I) stimulations. Comparison of the average strain amplitudes (J) and anisotropy coefficient A.C. (K) in the respective areas of stimulation. Data are presented as Tukey box plots with n = 17 microtissues. ****p < 0.0001. n.s. stands for non-significant (i.e. p > 0.05). Confocal projection of the fluorescent staining of actin in a representative microtissue (L), magnifications and color-coded map of the actin orientation (M) and polar plot (N) of the orientations of actin fibers in function of their localization, either in the center (red) or the left side (blue). Confocal projection of the fluorescent staining of collagen in a representative microtissue (O), magnifications and color-coded map of the collagen orientation (P) and polar plot (Q) of the orientations of collagen fibers in function of their localization, either in the center (red) or the left side (blue). Data of actin and collagen orientations are presented as mean ± SD with n = 16 microtissues and are significantly different (****p < 0.0001). Scale bar is 100 μm.

We then assessed the mechanical properties of microtissues 24 h and 48 h after seeding. We observed an almost doubling of the elastic modulus from 15.1 ± 6.5 kPa to 28.5 ± 11.1 kPa over these 24h, while the change in the delay *τ_S_* was not significant (Supp. Fig. 3). Since collagen was shown to be a key component of the tissue stiffness ^9^, we also assessed the rheological properties of microtissue composed of different collagen densities. We measured a very slight increase of stiffness from 15.1 ± 6.5 to 19.0 ± 7.0 kPa for initial collagen densities varying between 1.5 and 2.5 mg/mL, once again without any significant change of viscosity (Supp. Fig. 3). These values correlate with previous measurements ^9^ and the small variation of stiffness despite the large difference in initial collagen density is coherent with the insensitivity of stiffness to density for collagen under tension ^15^.

However, since collagen density feedbacks over time to regulate cell forces and stiffness ^9^, we made good use of our non-destructive approach to assess the possibly rapid changes of tissue stiffness and viscosity upon collagenase treatment. After 10 min of culture with 50 μg/mL of collagenase type I, we measured a strong decrease in tissue stiffness from 21.4 ± 8.3 to 9.1 ± 6.0 kPa, while the time delay between maximum stress and stretch was exactly the same before and after collagenase treatment. These results thus demonstrate that the cleavage of the peptide bonds of collagen by the collagenase strongly impacted the collagen structural stiffness without affecting the microtissue viscosity.

Finally, we explored the possibility to track changes in tissue mechanics during fibrogenesis. To this end, we induced the differentiation of fibroblasts to myofibroblasts by incubating microtissues with TGF-β1, a well-known, strong fribogenic growth factor ^9,31,32^. We first confirmed the differentiation of opto-RhoA fibroblasts into opto-RhoA myofibroblasts by observing a strong increase of α-SMA stress fibers after 3 days of treatment with TGF-β_1_ (Supp. Fig. 6.A-C). We then assessed the elastic modulus *E* and the delay *τS* by optogenetics after 24 h and 72 h of culture with or without TGF-β_1_. Similarly to previously reported measurements ^6,9,32^, we observed a strong increase of tissue stiffness for TGF-β_1_-treated microtissues at day 3 (Supp. Fig. 6.D). We also observed that the delay *τS* between maximum stress and stretch is almost twice as long with versus without TGF-β_1_ after 72 h (Supp. Fig. 6.E), which either supports a mechanism of stretch-induced perturbation of the cytoskeleton similar to the one we observed when varying the cantilever spring constant, or suggests a different relaxation dynamic of myofibroblasts versus fibroblasts.

Altogether, these results confirm the major role of collagen structure in the global stiffness of microtissues, whereas microtissue viscosity is mostly regulated by the contractile prestress of cells. As the collagen structure depends on the microtissue geometry ^25^, we next sought to probe local mechanical anisotropies using our optogenetic approach.

### Optogenetic assessment of local anisotropies

As microtissues suspended between two cantilevers exhibit a pronounced, homogeneous anisotropy along the x-axis (Fig. 2), we generated microtissues suspended between four cantilevers, in order to induce a more heterogeneous architecture presenting various degrees of anisotropy ^25^. We then performed a discoidal, isotropic light stimulation either in the center or on the left side of these square microtissues (Fig. 5.A-B, Supp. movie 6-7). We thus observed a completely isotropic displacement field for the centered stimulation, with a mean angle of 44 ± 4 °, whereas the eccentric stimulation induced a slightly biased displacement field with a mean angle of 53 ± 4 ° and more than 45% of the displacements oriented within less than 30 ° of the y-axis (Fig. 5. C-E, Supp. movie 6-7). These fields of displacement were paralleled with either an isotropic strain field for the centered stimulation, i.e. *ε_xx_ = ε_yy_* = −1.6 ± 0.2 %, or an anisotropic strain field for the eccentric stimulation with *ε_xx_* = −0.9 ± 0.2 % while *ε_yy_* = −1.6 ± 0.3 % (Fig. 2.F-J, Supp. movie 6-7). We thus obtained an A.C. of 0.0 ± 0.1 for the center part of the tissue whereas its left side exhibited an A.C. of −0.3 ± 0.1, indicating a moderate anisotropy along the y-axis (Fig. 5.K), similarly to previous results obtained utilizing finite element models ^25^. We then sought to compare these local anisotropic properties to the structural organization of these square microtissues. We simultaneously stained actin and collagen and quantified their respective orientations (Fig. 5.L-Q, Supp. movie 8). In good agreement with previous studies ^22,25^, we found a mostly random organization of actin and collagen fibers in the center of the microtissues, while 65 % of the actin fibers and 56 % of the collagen aligned within 30° of the y-axis along the left side. Bose et al. previously demonstrated that both ECM fiber alignment and density, resulting from the history of tissue formation, influence the local tissue stiffness ^25^. The correlation between the strain pattern, probed via optogenetics, the fine-scale cytoskeletal and extracellular architecture, and possibly stiffness heterogeneities thus demonstrates that the spatial propagation of mechanical signals in microtissues is strongly dependent upon actin and collagen organization, which in turn depend on the formation history of the tissue.

Finally, we investigated whether optogenetics could also be used to influence this architecture and the resulting local contractility. We conditioned the top half of a square microtissue with repetitive stimulations every 5 minutes for the first 24 h of its formation, before probing its contractility with a single, narrow, perpendicular pulse (Supp. Fig. 7.A-B). We thus observed a larger compression of the conditioned top half, reaching 3.0 ± 0.5 %, compared to a 2.1 ± 0.7 % compression of the non-conditioned bottom half (Supp. Fig. 7.C-E). The top half of the microtissues also exhibited a slightly higher signal of actin fluorescence (Supp. Fig. 7.F-H) but no difference in collagen fluorescence (Supp. Fig. 7.I-K) after immuno-staining. Altogether, these results suggest that, similarly to long term, external mechanical stimulations, a long term optogenetic conditioning during tissue formation impacts the expression and/or maturation of the actomyosin machinery ^11,33^, with the additional ability to do it locally.

Consequently, the use of optogenetically-modified cells as actuators to probe their environment appears as a powerful tool to guide tissue formation while simultaneously probing non-destructively architectural heterogeneities.

## Discussion

The mechanical properties of microtissues have been identified as major factors that regulate the physical integrity, contractility and function of such microtissues ^3–5,9^. Yet, these properties have only been examined through the application of external loading and biochemical treatments ^6,11,32^, which yielded key insights into how cells and ECM architecture impact microtissues’ mechanics. However, external actuations are inherently different than cell-generated forces, and the low spatial and temporal resolution of biochemicals limits their use for studying how cells probe the mechanics of their environment and how these mechanical perturbations propagate within the surrounding tissue. By introducing optogenetic modulation of the cellular contractility to the microtissues, we were able to produce local, physiological, cell-generated mechanical perturbations while simultaneously tracking their spatial and temporal propagation. These perturbations were reversible and paralleled by a compression of the stimulated zone, while non-stimulated zones were stretched.

We thus used optogenetic cells as biological actuators to probe the mechanics of fibroblasts/collagen microtissues. By tracking the maximum amplitudes of stress, compression and stretch in response to local light-activation of the contractility, we were able to plot hysteretic stress/strain curves, as the strains were delayed in time compared to the stress. We first observed that these fibrous microtissues seemed more compliant in compression than in stretch, as the compressed, stimulated part was more deformed than the stretched, non-stimulated one. Previous work has shown the highly nonlinear behavior of collagen gels, with a lower stiffness in compression compared to stretch ^34,35^, and an appropriate modeling of our light-sensitive microtissues would be particularly relevant to characterize this mechanical behavior in more details. We further demonstrated that, within the range of stimulation area and intensity tested in this work, single light-induced contractions did not induce plastic deformation of the microtissue. However, the amplitude of the contraction, modulated via the light-stimulation or the differentiation of fibroblasts into myofibroblasts, impacted the delay *τ_S_* between stress and strain. These results thus evidence a stress-dependent dynamic of strain propagation in fibrous microtissues, similarly to previous work in fibroblast populated collagen matrices that described an increase of the loss modulus with the strain rate ^26^ or an increase in stress-strain hysteresis with the strain rate ^36^, or in presence of myofibroblasts ^32^. One hypothesis for this behavior is a dependence between amplitude and dynamics of stress. However, we did not measure any difference in the time from zero to maximum contraction stress when varying the light intensity or the cantilever spring constant. A second hypothesis is an inherent stress-dependent viscosity of the collagen matrix but the inexistent difference in *τ_S_* when the collagen density is varied or when the collagen is cleaved with collagenase proves otherwise. A third hypothesis is the catch bond behavior of the contractile cytoskeleton that may induce such stress-dependent strain propagation. Although the catch-bond behavior of actin-myosin or cadherin-catenin bonds was demonstrated in single molecules ^27,29^ or single cells experiments ^28^, our results motivate further experimental studies to test whether this hypothesis is indeed correct in fibrous tissues, possibly via the use of cytoskeleton-associated tension reporters ^37–39^.

Using this optogenetic approach, we found that boundary conditions affect the stiffness of microtissues and, because of their impact on cell-generated stress ^2,9^, the delay between stress and strain, whereas collagen density or its degradation by collagenase only impact tissue stiffness. In the long term, we showed that the tissue stiffness increases over maturation time, especially in the case of a fibrogenic maturation, which correlates well with a cell-driven cross-linking of the matrix during tissue maturation or fibrosis ^32,40,41^.

These results are in good agreement with previous work showing that, in the short term, the collagen matrix is the main determinant of the tissue stiffness ^9,11^ while the actomyosin cytoskeleton is predominantly responsible for the viscoelasticity of microtissues ^13,42^.

Furthermore, we showed that the orientations and magnitudes of the light-induced contractions correlated with the collagen and actin orientation, which is known to influence in turn local tissue stiffness ^25^, and that repetitive stimulations during tissue formation could influence the expression and/or maturation of the actomyosin cytoskeleton, highlighting the predominant role of fibrillar architecture in the spatial propagation of mechanical signals. As collagen organization has been shown to be key to the long-scale interactions between cells and their navigation through the ECM ^43–45^, a more precise characterization using second-harmonic microscopy would be key to delineate the respective roles of thin isolated fibers and thick, strongly anisotropic collagen bundles.

Moreover, although we observed a correlation between cell-induced strain patterns and tissue architecture, we cannot deduce from our experiment the cause of the anisotropic strain pattern, whether it is due to the cytoskeletal and extracellular organization, the spatial heterogeneity in stiffness or a combination of both parameters. Microtissue regions with aligned actin and collagen fibers were shown to be significantly stiffer than randomly organized regions ^25^, but unraveling the respective roles of the different types of anisotropy (i.e. anisotropy of cellular contraction, of mechanical properties, of fiber orientations) remains currently an extremely complex task that has only begun to be undertaken, and would require further experimental studies, possibly combining tissue engineering, optogenetics, cytoskeleton-associated tension reporters and computational modeling ^22,25,37–39^. Nevertheless, the ability of our method to apply dynamic, internal mechanical perturbations, while simultaneously measuring the magnitude and spatio-temporal propagation of such perturbations opens an exciting avenue for guiding the formation of tissues while simultaneously assessing their architectural heterogeneities, as well as for examining the mechanical guidance of cell migration in fibrillar matrices.

However, as our approach uses the contraction of one part of the tissue (the stimulated one) to stretch the other part (the non-stimulated one) to probe the mechanical properties of the latter, it inherently requires an active actomyosin machinery, which could complicate the study of the role of cytoskeletal components in regulating tissue mechanics. Indeed, a complete inhibition of the contractile ability of the microtissue, using high doses of actomyosin-targeting drugs for example, would hinder the use of our approach. Yet, our method remains valid as long as the contractile machinery is at least partially active, e.g. for small to medium doses of actomyosin-targeting drugs. Similarly, although the recruitment of CRY2 to the membrane is very fast (few seconds), its dissociation is slower (several minutes) ^21^, while the dynamic of the RhoA pathway leads to a contractility activation in the order of tens of seconds and a slower relaxation of several minutes ^19^. As a result, the induced cell contraction and relaxation takes almost 20 minutes, thus precluding its use for probing rapid mechanical changes or quantifying time-dependent mechanical responses of microtissues at different frequencies. Finally, our optogenetic target RhoA has many downstream effectors. Although we used single, short stimulations that induced an elastic response of the microtissue with no long lasting effects such as cytoskeleton remodeling for example, we cannot exclude that off target effects could exist, via for example the self-amplification and self-inhibition of RhoA ^46^.

In conclusion, the combination of tissue engineering and optogenetics provides unique opportunities to quantitatively demonstrate the impact of physical and biological parameters on the generation, propagation and sensing of cell-generated mechanical perturbations in 3D tissues. Most importantly, our approach paves the way to probing the rheology of 3D tissues in real time and non-destructively, using their own constituting cells as internal actuators. These same attributes will likely provide valuable opportunities to elucidate how mechanical cues dynamically regulate tissue formation and function over space and time. Altogether, our work thus demonstrates that an optogenetic control of cell contractility, combined with specifically designed microtissue geometry and possibly computational modeling ^25^, could offer a powerful approach to analyze the complex interplay between mechanical boundary constraints, cell contractility, ECM density, alignment, and mechanical properties.

## Materials & Methods

### Cell culture, reagents and immuno-stainings

The DHPH domain of ARHGEF11 Guanine Exchange Factor gene was cloned into CRY2PHRmCherry using Nhe1 and Xho1 cloning sites. ArhGEF11-CRY2PHR-mCherry and CIBN-GFP-CAAX were then inserted into lentiviral backbones (pHR and pLVX respectively) to create the stable cell line of opto-RhoA fibroblasts from NIH 3T3 fibroblasts (ATCC). After viral transduction, only the top 2% of the cells presenting the highest expression levels for both transcends were FACS-sorted before amplification. The obtained opto-RhoA fibroblasts (< 15 passages, kindly provided by L. Valon and M. Coppey, Institute Curie, Paris, France) were cultured in Dulbecco’s modified Eagle’s medium (DMEM, Gibco Invitrogen) supplemented with 10% fetal bovine serum (FBS, Gibco Invitrogen), 100U/ml of penicillin and 100 μg/ml of streptomycin (Gibco Invitrogen), and kept at 37°C in an atmosphere saturated in humidity and containing 5% CO2. Collagenase experiments were performed by incubating microtissues with 0.05 g/L of collagenase type I (Sigma) for 10 minutes before thorough rinsing with PBS and replacement with growth medium. To induce myofibroblasts differentiation in the microtissues, regular culture media was supplemented with 5 ng/mL of TGF-β1, (T7039, Sigma) just after cell seeding. Prior to immunostaining, samples were fixed in 4% paraformaldehyde (Sigma) and blocked with 2% BSA Sigma). Collagen and α-smooth muscle actin (α-SMA) were immuno-stained with a primary antibody against Collagen type I (Sigma) and α-SMA (Sigma), respectively, and detected with Alexa 647-conjugated, isotype-specific, anti-IgG antibodies (Invitrogen). Samples were permeabilized with 0.5 % Triton X-100 (Sigma) in TBS (50 mM Tris-HCl, 0.15 M NaCl, pH 7.4) either before or after the incubation with the primary antibody, in order to stain for intracellular α-SMA or extracellular collagen I, respectively. Actin was labeled with phalloidin-Atto488 (Sigma) and nuclei with DAPI (Invitrogen).

### Device fabrication and calibration

The microtissues are engineered in microwells containing two or four T-shape cantilevers. This T-shape is essential to constrain the microtissue formation, ensure good anchorage of the microtissue to its supporting cantilevers and avoid its slipping from the post. To achieve this complex geometry with a top cap wider than its post, SU-8-based masters were fabricated following the technique described previously ^2,47^. Briefly, successive layers of negative and positive photoresist (Microchem) were spin coated, insolated and baked to create multilayers templates. A first layer of negative resist allows the creation of the posts, a second one consisting of a mix of 70 % negative and 30 % positive photoresist serves as a lithographic-stop layer that prevents unwanted cross-linking of the underlying layer, and a third layer of negative photoresist leads to the creation of the top wide cap at the tip of the post. Polydimethylsiloxane (PDMS, Sylgard 184, Dow-Corning) microfabricated tissue gauges (μTUGs) were molded from the SU-8-based masters by double replication as described previously ^2,47^. Briefly, SU-8-based masters or PDMS replicates were oxidized in an air plasma (REF), silanized with trichloro(1H,1H,2H,2H-perfluorooctyl)silane (Sigma) vapor overnight under vacuum to facilitate subsequent release of PDMS from the template and avoid tearing the top cap of the cantilevers. Prepolymer of PDMS was then poured over the template, degassed under vacuum, cured at 65°C for 20 h, and peeled off the template. Different ratios of PDMS/curing agents were used to modulate PDMS stiffness, which was assessed through uniaxial extension of 50×4×1 mm strips with an Instron 5848 Microtester (Instron). Stiffness was determined from the linear region of the obtained stress-strain curves. Cantilever spring constants were calibrated with a capacitive MEMS force sensor mounted on a micromanipulator as described previously ^4,22^. Briefly, the sensor tip was placed 20 μm below the top of the cap and the probe translated laterally against the outer edge of the cantilever. The spring constant was calculated from the displacement of the cantilever head and the reported sensor force. The spring constant of the cantilevers were found to be 0.20 ± 0.03 N/m, 0.45 ± 0.10 N/m and 1.10 ± 0.26 N/m for PDMS/curing agent of 1:20, 1:10 and 1:4, respectively. The dimensions and geometry of the cantilevers were regularly assessed to ensure they were not affected by successive replications.

Before cell seeding, the PDMS templates were sterilized in 70% ethanol followed by UV irradiation for 15 min and treated with 0.2% Pluronic F127 (Sigma) for 2 min to reduce cell adhesion. A reconstitution mixture, consisting of 1.5 mg/mL or 2.5 mg/mL liquid neutralized collagen I from rat tail (Advanced Biomatrix) was then added to the surface of the substrates on ice and templates were degassed under vacuum to remove bubbles in the liquid. A cooled suspension of 750,000 cells within reconstitution mixture was then added to the substrate and the entire assembly was centrifuged to drive the cells into the micropatterned wells, resulting in approximately 500 cells per well. Excess collagen and cells were removed by de-wetting the surface of the substrate before incubating at 37°C to induce collagen polymerization for 9 min. Culture medium was then added to each substrate. Microtissues were kept in the incubator for 24h before stimulation experiments, unless specified otherwise. Over these first 24 h of cultivation, the fibroblasts spread and spontaneously compact the collagen matrix (Fig. 1). The two T-shaped cantilevers anchor and constrain the contraction of the collagen matrix to form a microtissue that spans across the top of the pair of cantilevers.

### Optogenetic stimulation and microscopy

Optogenetics stimulation and brightfield imaging were performed using an inverted Nikon Eclipse TI-2 microscope with an Orca flash 4.0 LT digital CMOS camera (Hamamatsu) and a CFI S Plan Fluor ELWD 20x/0.45 objective (Nikon). Light stimulations were achieved using a Mosaïc 3 digital micromirror device (DMD, Andor) and the 470 nm diode of a Spectra X light source (Lumencor). The light power was calibrated using a photodiode power sensor S120C (Thorlabs), placed in the light path, after the objective, and connected to a compact power and energy meter console PM100D (Thorlabs). The total irradiance sent to the microtissue could be varied from 0 to 4.8 mW/mm^2^. Microtissues were stimulated with an irradiance of 1.6 mW/mm^2^ for 500 ms for all the experiments, unless specified otherwise. The light pattern, intensity and duration, as well as the microscope itself, were controlled using NIS Elements software (Nikon). Microtissues were maintained at 37 °C and 5% CO2 in a top-stage incubator (Oko Lab).

Confocal images were obtained with a Leica laser scanning microscope (LSM SP8, Leica) equipped with a plan apochromatic 40x/1.30 objective (Leica). Actin and collagen orientations were evaluated from confocal z-stack images with the Orientation-J plug-in (http://bigwww.epfl.ch/demo/orientationj/) ^48^ in Image J. This plugin computes the structure tensor for each pixel in the image by sliding a Gaussian analysis window (variance σ = 2 pixels) over the entire image. From the structure tensor are extracted both the orientation and coherency properties of the region of interest. These properties are then gathered in a color map in HSB (Hue Saturation Brightness) mode where the hue corresponds to the orientation, the saturation to the coherency and the brightness to the source image. The coherency indicates if the local image features are coherently oriented or not. The polar histograms presented in Fig. 2 and Fig. 5 are weighted histograms, the weight being the coherency. Consequently, thicker or denser collagen bundles are weighted more than isolated, randomly oriented fibers, in coherence with the fact that thick and dense collagen bundles weight more in the mechanical behavior of a fibrous microtissue than thin random fibers ^25^.

### Force measurement and strain measurement

The contraction force generated by individual microtissues before, during and after light stimulations was assessed from the deflection of the cantilevers. This deflection *d* was determined by comparing the position of the top of the cantilevers to their initial position (i.e. before seeding the cell/collagen mixture). To this end, brightfield images were taken every 30 s and the displacement of the top of the cantilevers was tracked using custom MATLAB script. Briefly, the displacement of the sharp contrast created by the edge of the cantilever heads was tracked over time by auto-correlation of the interpolated line profiles, allowing subpixel resolution. Tracking results were visually checked and faulty tracking discarded. Based on the linear bending theory, the resulting force *F* applied on a cantilever was inferred from its deflection and its spring constant *k* as follow: *F* = *k*.*d*. The force generated by one microtissue corresponds to the average force applied to each of its two anchoring cantilevers. Only tissues that were uniformly anchored to the tips of both cantilevers were included in the analysis. This anchorage between microtissue and cantilevers was visually checked over the whole duration of the experiment. Only microtissues wrapping completely the cap of both cantilevers were selected (Supp. Fig. 8.A-C). Tissues tearing or slipping from their cantilevers during an experiment were discarded from the analysis.

The stress *σ_xx_* was calculated by dividing the force by the cross-sectional area measured in the center of the tissue. The width of each microtissue (i.e. its dimension along the y-axis, the x-axis being between the two cantilevers) was tracked during stimulation experiments. The cross-sectional areas were measured from confocal z-stack images of tissues fixed and stained right after experiments, in their center, similarly to previous works (Supp. Fig. 8.D-H) ^2,4,6,9,11,25^. No measurable change in cross-sectional area was observed over the duration of light-stimulation experiments, in agreement with the almost inexistent displacements along the y-axis (Fig. 1–4). Consequently, the cross-sectional areas were considered unchanged over the duration of stimulation. Of note, the cross-sectional areas were linearly correlated with the tissue width squared *w*^2^ (Supp. Fig. 8.I-J, R^2^ = 0.80), which allowed to estimate tissue cross-section when microtissues could not be fixed immediately after experiments. The displacement fields were determined from the brightfield images using a particle image velocimetry (PIV) algorithm implemented as a Matlab toolbox (https://pivlab.blogspot.com/) ^49^. Briefly, small sub images (interrogation areas) of an image pair consisting of the reference image at t = 0 and the image of interest were cross-correlated in the frequency domain using FFT to derive the most probable particle displacement vector in the interrogation areas, with its *x* and *y* components *u* and *v*, respectively. Analyzed images were 1024×512 pixels (800 × 400 μm) or 1024×1024 pixels (800×800 μm) for microtissues between 2 or 4 cantilevers, respectively, and the size of the interrogation areas was successively reduced from 128×x128 to 96×96 and finally 64×64 pixels (corresponding to 100×100, 75×75 and 25×25 μm, respectively). For each pass, i.e. for each size of interrogation area, the overlap between interrogation areas was set to 50%, leading to a final resolution of 25 μm. Outliers were filtered using a local normalized median median filter ^50^, missing vectors were replaced by interpolated data ^51^ and the noise was reduced using a penalized least squares method ^52^.

The strain maps were derived by numerical differentiation of *u* and *v* to both *x* and *y*, comparing the displacement vector of each particle with the vectors of surrounding particles in a window of 3×3 particles ^53^:

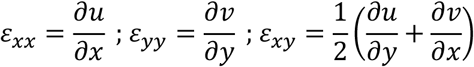

For plotting stretch and compression over time, the *ε_xx_* strain was estimated by averaging the displacement *u* (i.e. along the x axis) over the width of the tissue before calculating its slope across the length of the stimulated and the non-stimulated areas, respectively. The tissue was considered as homogeneous and was not discretized as individual cells and matrix. Consequently, we measured tissue displacements and corresponding tissue deformations. Of note, cell migration over the duration of a contraction was considered negligible compared to cell deformation.

### Data analysis and statistics

For each line graph, the mean represents the mean ± standard deviation of *n* microtissues (*n* is defined in the caption of each figure). For each box plots, the box extends from the 25th to 75th percentiles, the median is plotted as a line inside the box and the whiskers extend to the most extreme data point that is no more than 1.5 times the interquartile range (IQR) from the edge of the box (Tukey style). Significances were assessed with Prism (GraphPad) by one-way analysis of variance, using Tukey’s significant difference test (p < 0.001 was considered significant).

## Data availability

The datasets generated during and/or analyzed during the current study are available from the corresponding author on reasonable request.

## Code availability

Matlab analysis procedures can be made available upon request to the corresponding author.

## Aknowledgments

A.M. acknowledges support from the University Grenoble Alpes Ph.D. fellowship program. G.C. acknowledges financial support from the ANR SupraWaves project, grant ANR-19-CE13-0028 of the French Agence Nationale de la Recherche (ANR). M.B. acknowledges financial support from the ANR MechanoSwitch project, grant ANR-17-CE30–0032-01. T.B. acknowledges fundings through CNRS grants (Actions Interdisciplinaires 2017, DEFI Instrumentation aux limites 2017, Tremplin@INP 2021, PEPS CNRS-INSIS 2021, Lumière Visible et Vie 2022). This work was supported by the Center of Excellence of Multifunctional Architectured Materials “CEMAM” (n° AN-10-LABX-44-01).

The authors thank P. Moreau and I. Wang for their technical support, L. Vallon, S. de Beco and M. Coppey for kindly providing the opto-RhoA fibroblasts, G. Chagnon and N. Briot for the mechanical characterization of the PDMS, as well as P. Recho for helpful discussions.

## Supplemental Figures

**Supp. Figure 1.**
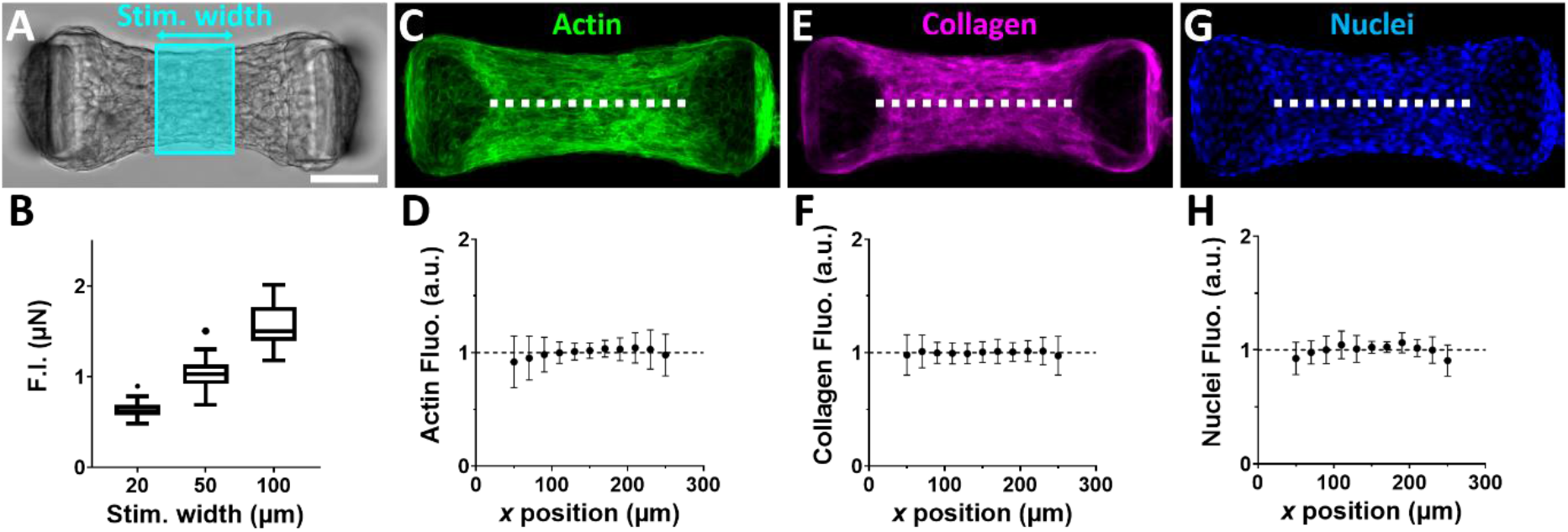
Light-induced contraction is proportional to the number of stimulated cells. Representative microtissue with the stimulation pattern in blue (A) and corresponding force increase (F.I.) in function of the stimulation width (B). Confocal projections and average fluorescence intensity along the dotted line for microtissues stained for actin (C, D), collagen (E, F) and nuclei (G, H). Data are presented as mean ± SD with n > 15 microtissues. Scale bar is 100 μm.

**Supp. Figure 2.**
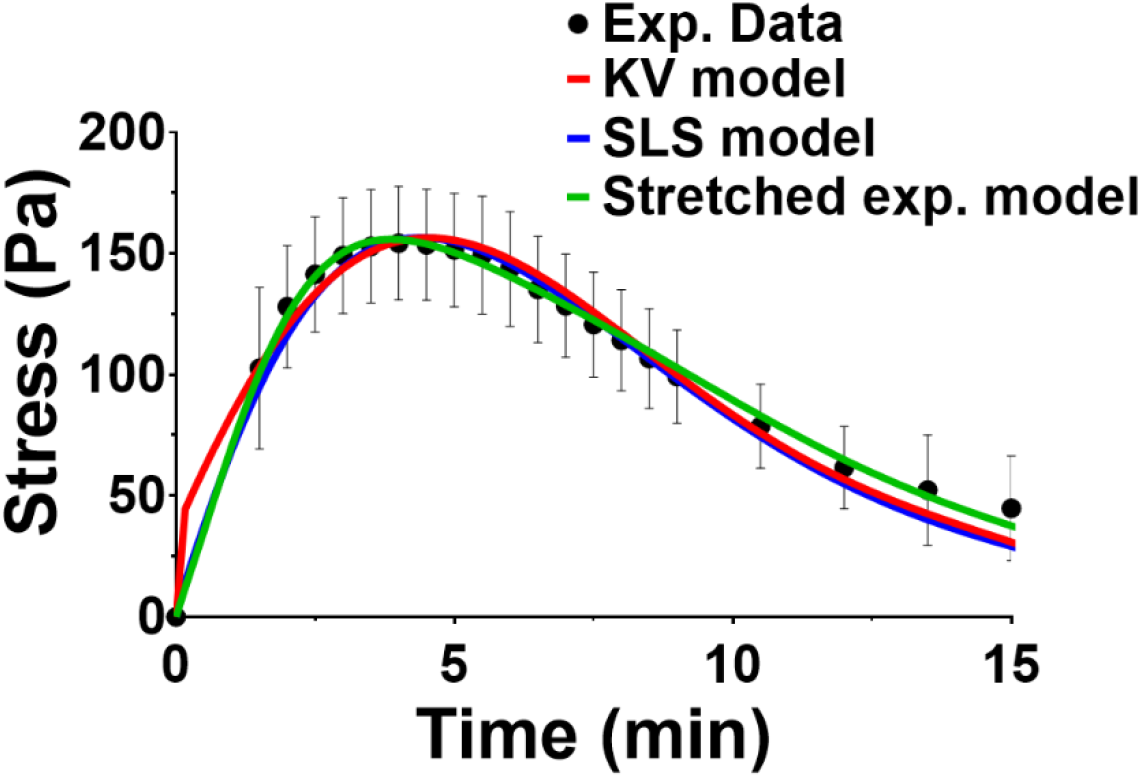
Viscoelastic modeling, based on ^13^, of stress response to a light stimulation at t = 0. The black dots correspond to the experimental data obtained in Figure 4 for a stimulation width of 50 μm, and presented as mean ± SD with n = 20 microtissues. The red curve corresponds to the Kelvin-Voigt (KV) model (a spring and a dashpot in parallel), the blue curve to the standard linear solid (SLS) model (a spring and a dashpot in series, in parallel to another spring), and the green curve to the stretched exponential model (a sum of Maxwell bodies, i.e. a spring and a dashpot in series, with a specific distribution of time constants). The corresponding equations and parameters are the following:

– KV model: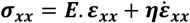 with *E* = 18.7 kPa and *η* = 1.9 MPa.s, leading to a retardation time 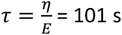.
– SLS model: 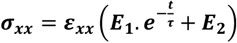 with *E_1_* = 56.3 kPa, *E_2_* = 14.7 kPa and *τ* = 135 s.
– Stretched exp. Model^13^: 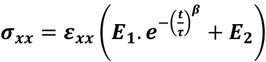 with *E_1_* = 65.0 kPa, *E_2_* = 10.0 kPa, *τ* = 129 s and *β* = 0.66

**Supp. Figure 3.**
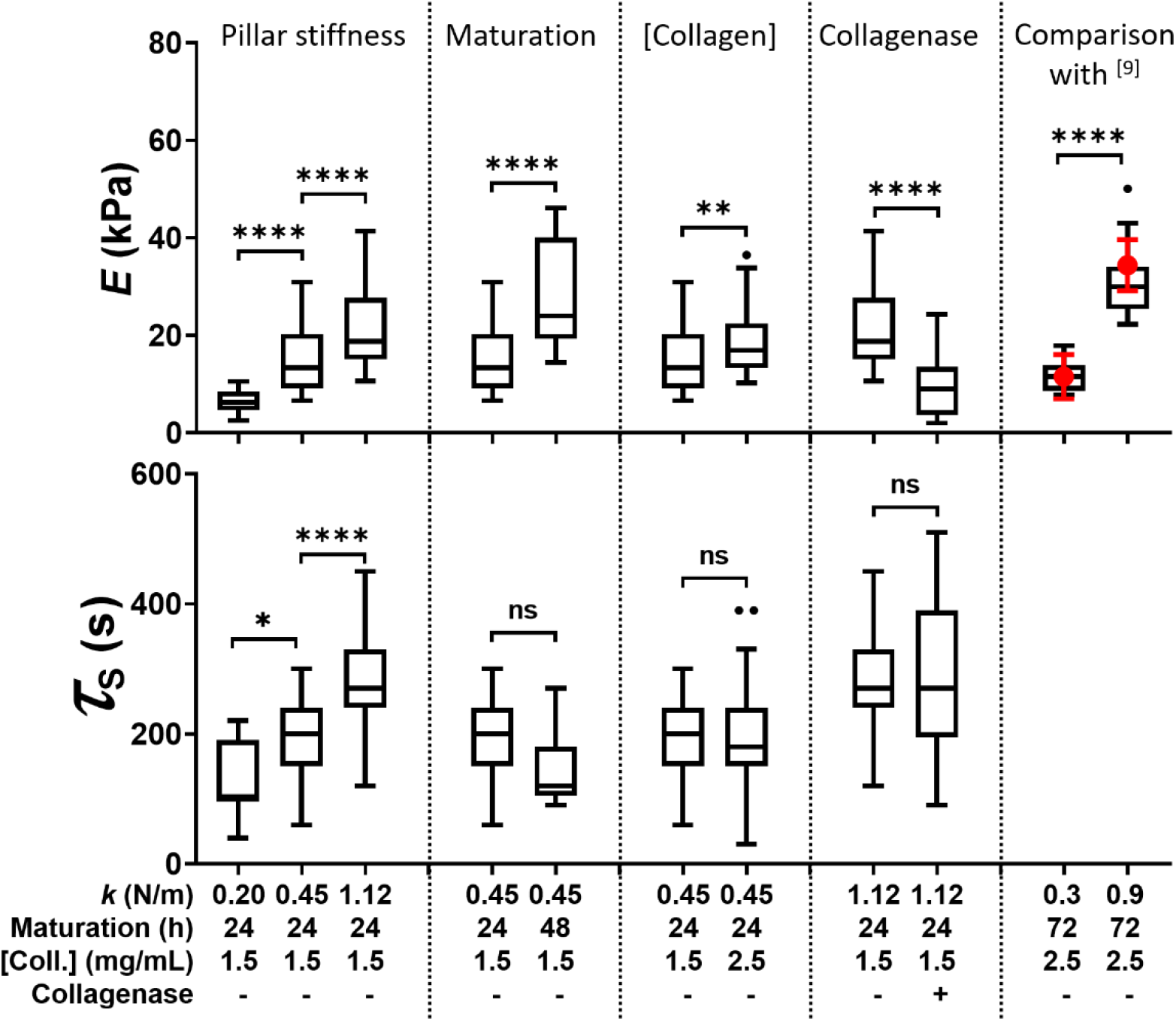
Optogenetic assessment of the rheological properties of microtissues. Elastic modulus *E* and delay *τ_S_* between maximum stress and stretch for three different spring constants *k* of the cantilevers, two durations of maturation, two collagen densities [Coll.] and the presence or absence of collagenase. Data are presented as Tukey box plots with n > 18 microtissues. The two last columns present elastic moduli obtained with this approach (box plots) and magnetic actuation in ^9^ (red dots) for the same culture conditions.

**Supp. Figure 4.**
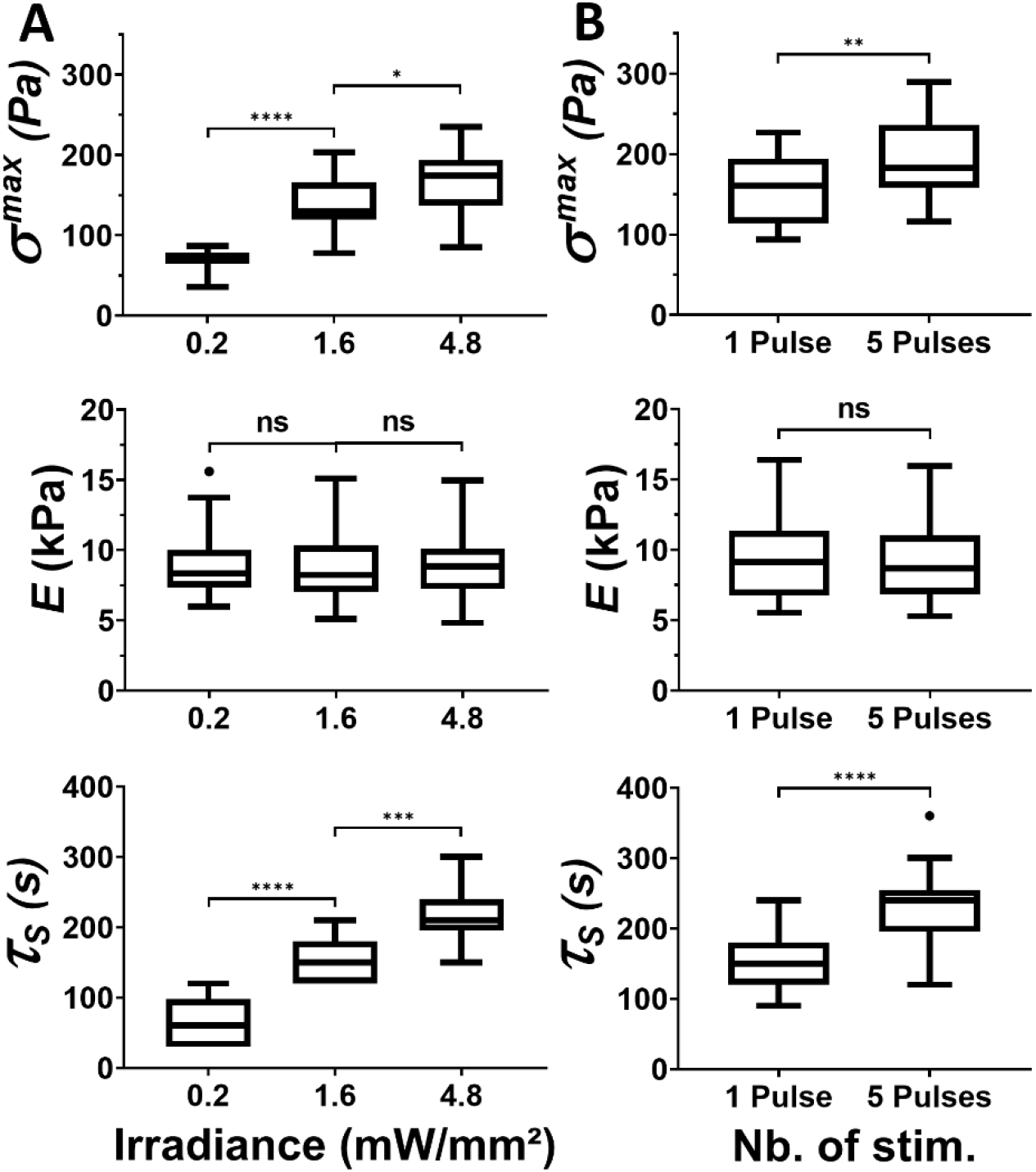
Influence of the amplitude of contraction on the measured mechanical properties. Light-induced maximum stress *σ ^max^*, elastic modulus *E* and delay *τ_S_* between maximum stress and *ε_xx_* strain for (A) an irradiance ranging from 0.2 to 4.8 mW/mm^2^ (n = 13 microtissues) and for (B) one pulse or a series of five, 1-minute-spaced pulses of 1.6 mW/mm^2^ (n = 20 microtissues). Data are presented as Tukey box plots.

**Supp. Figure 5.**
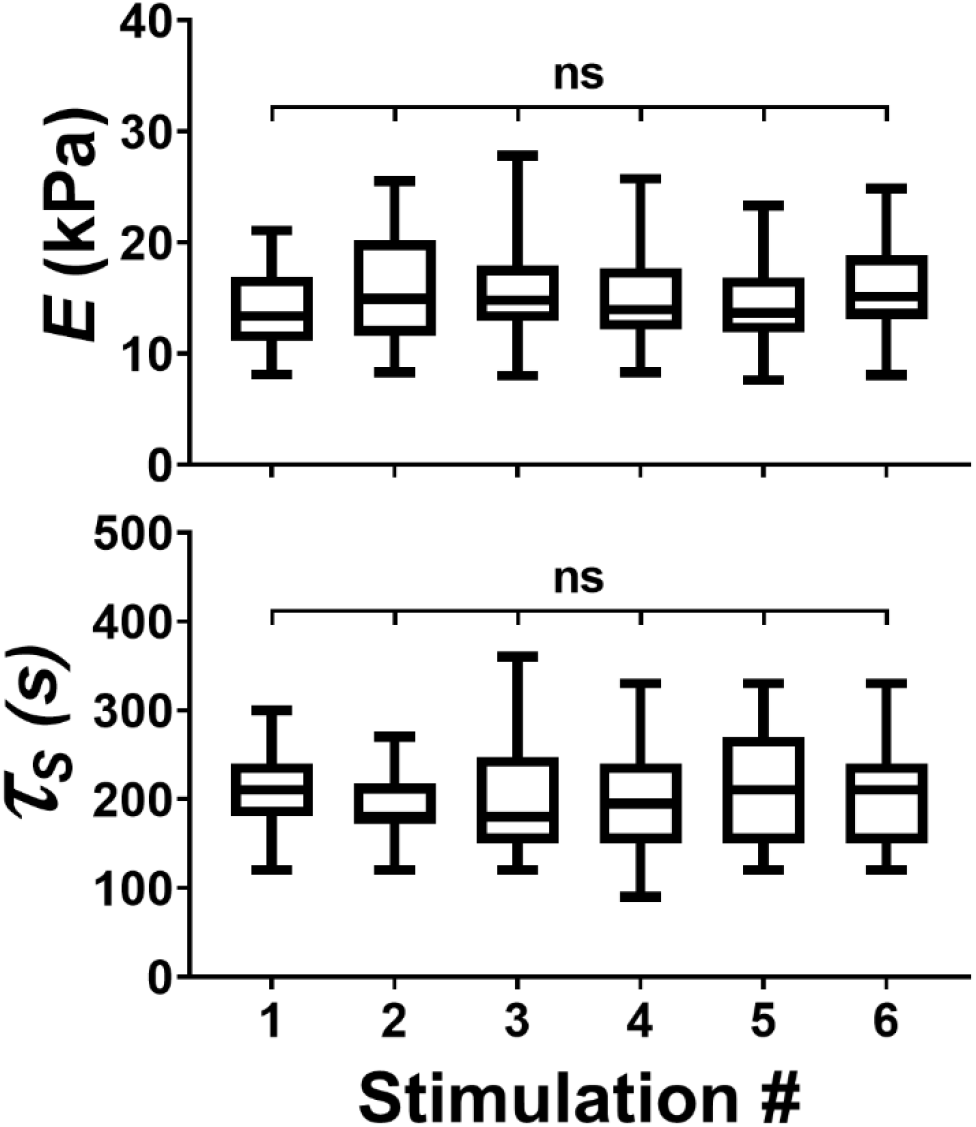
Optogenetic stimulations do not induce plastic, irreversible deformations of the microtissue. Elastic modulus *E* and delay *τ_S_* between maximum stress and *ε_xx_* stretch for six successive stimulations spaced far enough apart in time to allow complete relaxation (30 minutes between each stimulation). Data are presented as Tukey box plots with n > 29 microtissues.

**Supp. Figure 6.**
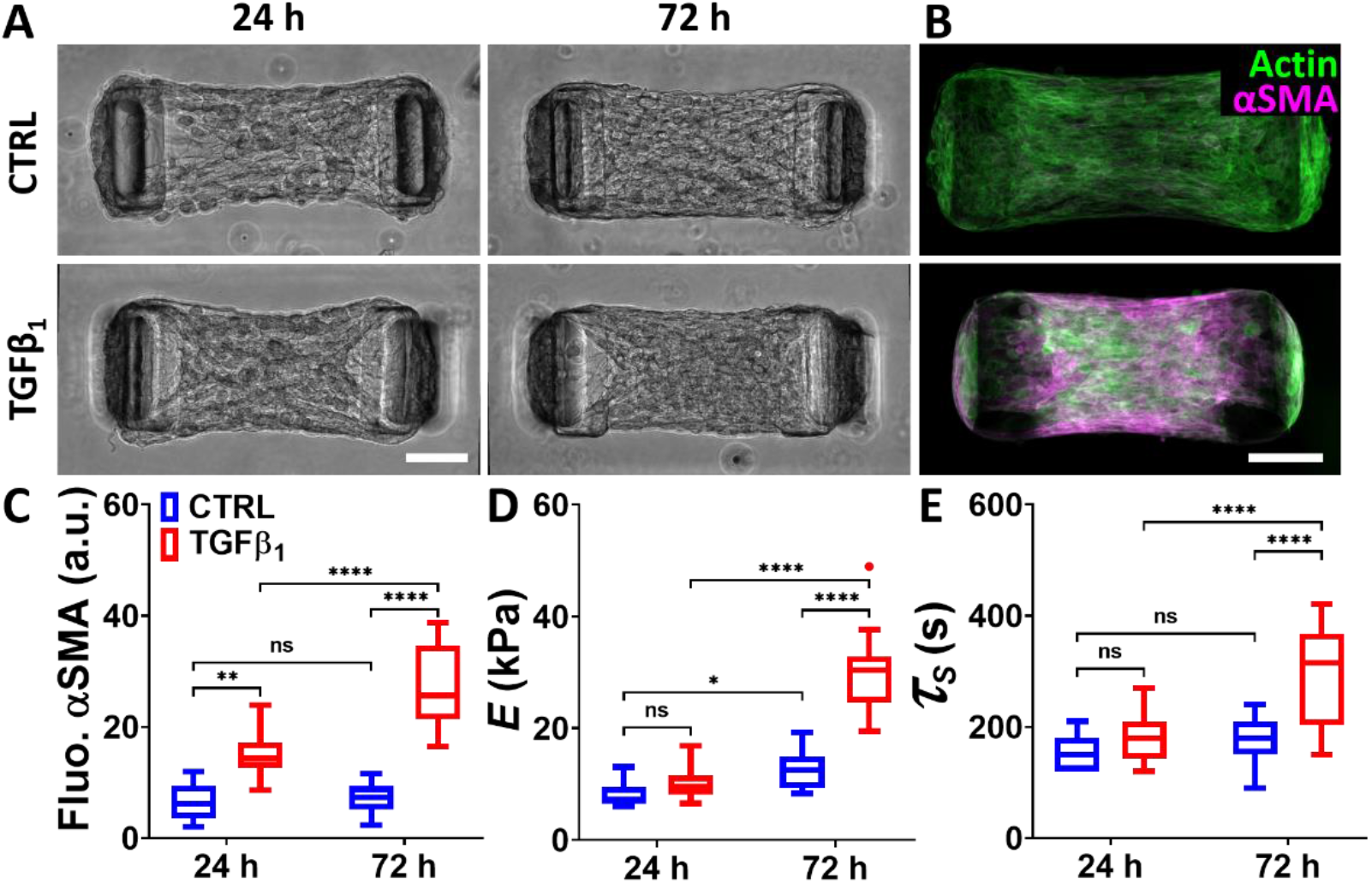
Myofibroblasts strongly impact the mechanical properties of microtissues. (A) Brightfield images of representative microtissues after 24 h and 72 h of culture in in serum-only (CTRL) and TGF-β1 treated medium. (B) Corresponding confocal projections for microtissues stained for actin (green) and α-SMA (magenta) at 72 h. Fluorescence intensity of the α-SMA staining (C), elastic modulus *E* (D) and delay *τ_S_* between maximum stress and *ε_xx_* strain (E) for microtissues after 24 h and 72 h of culture in in serum-only (CTRL, in blue) and TGF-β1 (in red) treated medium. Scale bars are 100 μm. Data are presented as Tukey box plots with n > 8 microtissues for (C) and n > 12 microtissues for (D) and (E).

**Supp. Figure 7.**
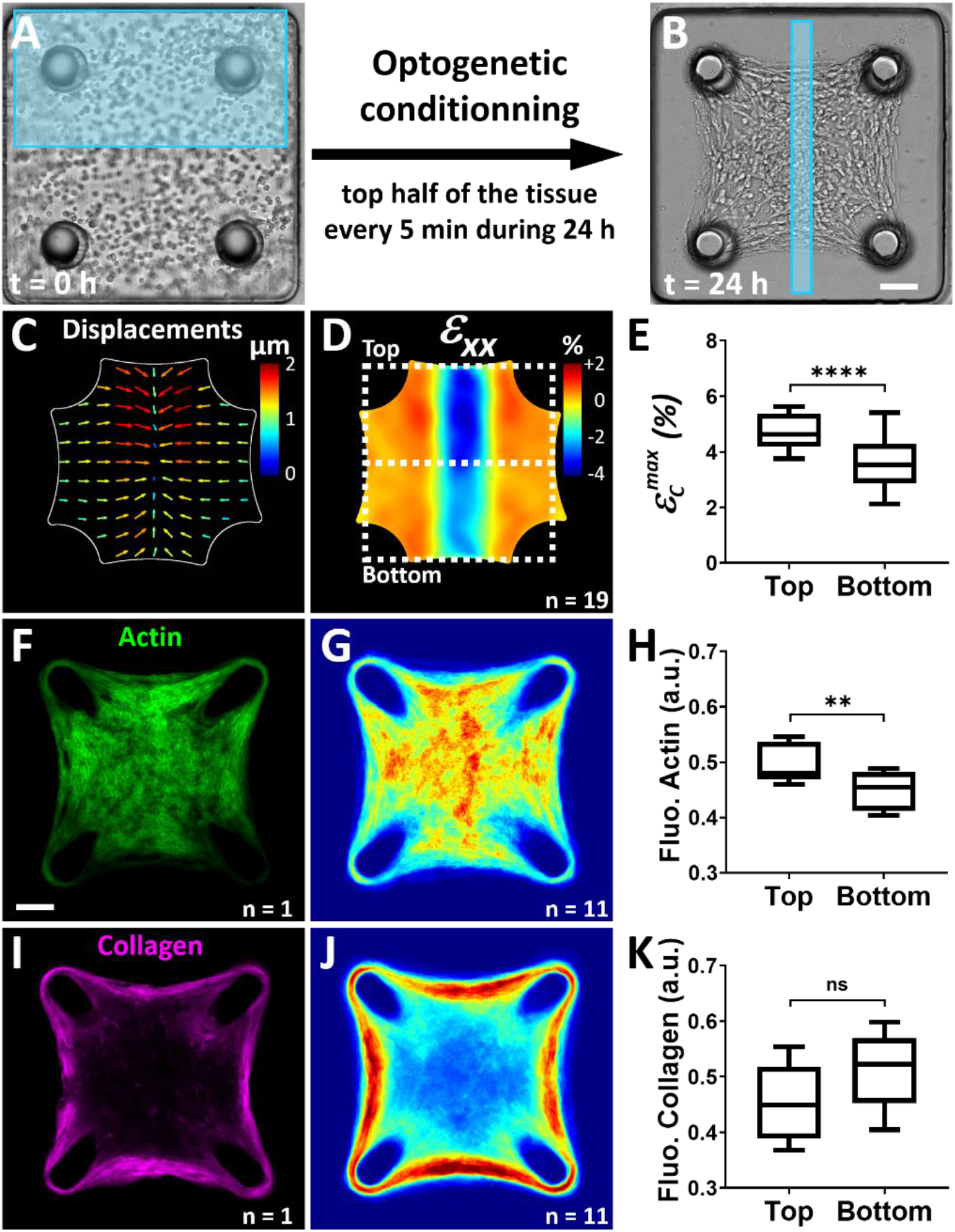
Optogenetic conditioning of microtissue contractility and architecture. (A) Representative square microtissue at t = 0 h. During the first 24 h of formation, the top half of the microtissue is stimulated every 5 min with a pulse of light (represented by the blue rectangle). (B) After 24 h of formation, a centered, rectangular area of the microtissue (in blue) is stimulated by a single light pulse. Resulting average displacement filed (C), *ε_xx_* strain field (D) and comparison of the maximum compression 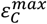 in the top, conditioned area and the bottom, non-conditioned area (E). Data are presented as Tukey box plots with n = 19 microtissues. ****p < 0.0001. Individual (F) and average (G) confocal projection of the fluorescent staining of actin. (H) Corresponding quantification of the fluorescence intensity of the actin staining in the top and bottom area. Individual (I) and average (J) confocal projection of the fluorescent staining of collagen. (K) Corresponding quantification of the fluorescence intensity of the collagen staining in the top and bottom area. Data of actin and collagen fluorescence are presented as Tukey box plots with n = 11 microtissues. **p < 0.01. n.s. stands for non-significant (i.e. p > 0.05). Scale bars are 100 μm.

**Supp. Figure 8.**
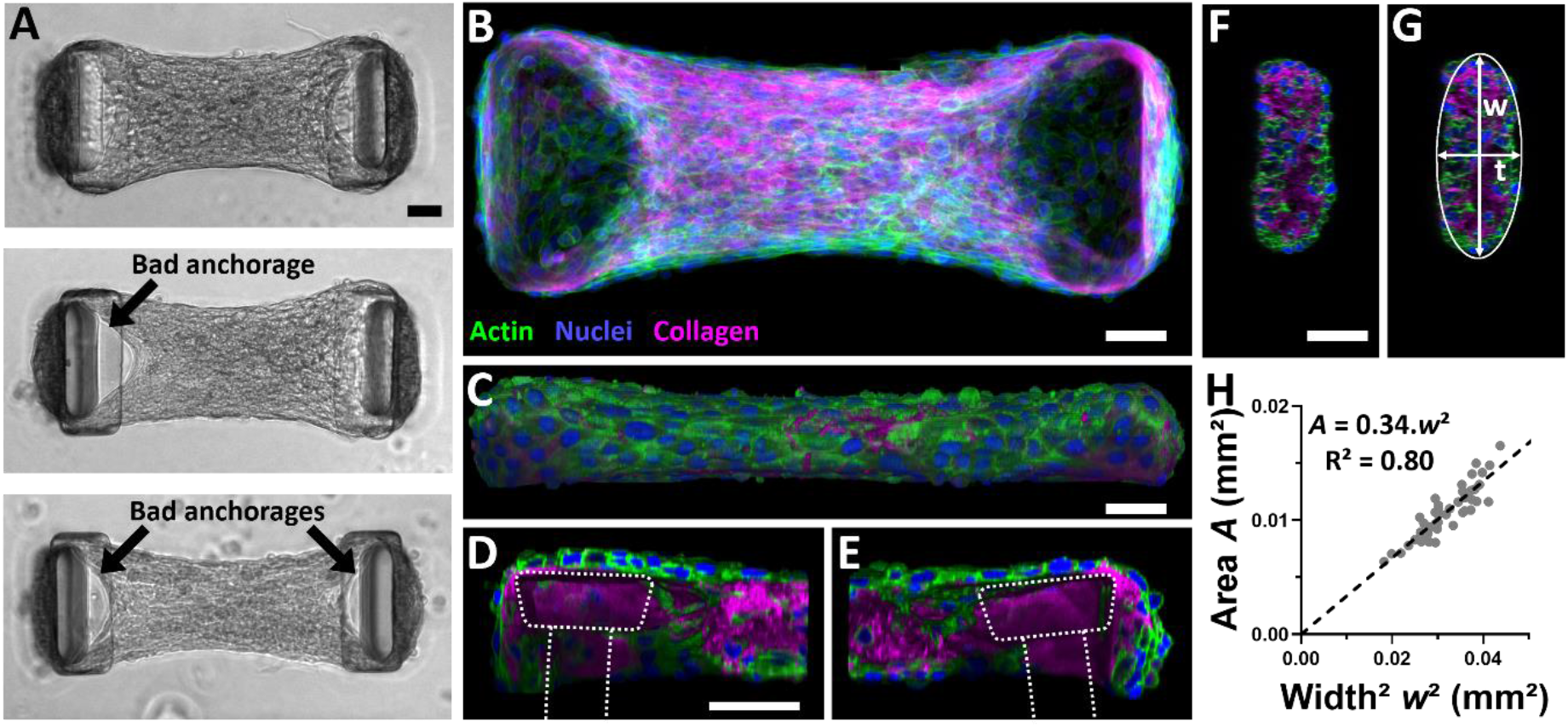
Anchorage of microtissues to cantilevers and measurement of the cross-section. (A) Representative microtissues properly anchored (top image), with cells wrapping completely the cap of both cantilevers, and poorly anchored to either one (middle image) or two (bottom image) of the cantilevers, with no cells on top of the cantilever and a tissue in a different focal plane than the cantilevers. Poorly anchored microtissues were not selected for experiments. (B) Representative top-view confocal projection of a microtissue stained for actin (green), nuclei (blue) and collagen (magenta). (C) Corresponding orthogonal view in the xz plane. Magnified orthogonal views of the left (D) and right (E) anchorages where cantilevers are outlined with dashed lines. (F) Orthogonal view in the yz plane for measuring tissue cross-section to derive tissue stress σxx. (G) The yz cross-section can be fitted with an ellipse of width w and thickness t. The ratio of the cross-sectional area to the squared width w^2^ was used to infer cross-sectional area (A = 0.34 w^2^, R^2^ = 0.80, n = 45 microtissues) from top view images when microtissues could not be fixed immediately after experiment. Scale bars are 50μm.

## Supplemental Movies

**Supp. Movie 1. Light-controlled local contraction of microtissues.** Time lapse of a representative opto-microtissue whose left-then right-half are stimulated with blue light (represented by a blue rectangle) at t = 40 s and t = 21 min, respectively. Upon blue light illumination, the microtissue contracts locally before slowly relaxing, as shown by the PIV-tracking of the displacements.

**Supp. Movie 2. Anisotropic contraction of an opto-microtissue upon an isotropic stimulation**. Temporal evolution of a representative opto-microtissue and the corresponding strain fields upon the stimulation in its center with a 50 μm diameter discoidal light pattern (represented by a blue disc) at t = 1’20”. Despite an isotropic stimulation, the resulting strain is dominated by its x-axis component, as shown by the temporal evolution of *ε_xx_* and *ε_yy_*.

**Supp. Movie 3. Confocal reconstruction of an opto-microtissue.** Z-stack (top) and 3D view (bottom) of a representative opto-microtissue stained for actin (in green), collagen (in magenta) and nuclei (in blue), highlighting the anisotropic orientation of actin and collagen fibers along the x-axis. Cantilevers are reconstructed from thresholded brightfield images.

**Supp. Movie 4. Light-induced local contractions evidence the viscoelastic properties of microtissues.** Temporal evolution of a representative opto-microtissue, displacement field and *ε_xx_* strain field upon the stimulation of its left-half with blue light (represented by a blue rectangle) at t = 2’30”. The stimulated half is strongly compressed, with a maximum at t = 7’30", while the non-stimulated half is stretched, with a maximum at t = 9’30”.

**Supp. Movie 5. Contraction is proportional to the area of stimulation.** Temporal evolution of a representative opto-microtissue, displacement field and *ε_xx_* strain field upon the illumination of its center by a 20 μm, a 50 μm and a 100 μm wide stimulation (represented by blue rectangles) at t = 2 min, t = 22 min and t = 42 min, respectively.

**Supp. Movie 6. Isotropic contraction of the center of an opto-microtissue.** Temporal evolution of a representative opto-microtissue, displacement field and strain fields upon the stimulation in its center with a 200 μm diameter discoidal light pattern (represented by a blue disc) at t = 40 s. The resulting contraction is isotropic, as shown by the displacement field and the similar amplitudes of *ε_xx_* and *ε_yy_*.

**Supp. Movie 7. Anisotropic contraction of the side of an opto-microtissue.** Temporal evolution of a representative opto-microtissue, displacement field and strain fields upon the stimulation on its left side with a 200 μm diameter discoidal light pattern (represented by a blue disc) at t = 40 s. The resulting contraction is anisotropic, as shown by the displacement field and the differences in amplitude between *ε_xx_* and *ε_yy_*.

**Supp. Movie 8. Confocal reconstruction of a square opto-microtissue.** Merged Z-stack (top left) and 3D view (top right) of a representative opto-microtissue stained for actin (in green) and collagen (in magenta). Separated Z-stacks (middle) and magnifications (bottom) of the actin (in green) and the collagen (in magenta) in the left and the center area, highlighting the anisotropic orientation of actin and collagen fibers along the sides, while the center is mostly disorganized. Of note, the intensity of the bottom magnifications is gamma corrected 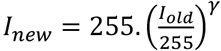, with *I_new_* and *I_old_* the intensity after and before correction, respectively, and *γ* = 0.6) to better visualize thin, dim collagen fibers despite the intense fluorescence of large collagen bundles.

